# DiCE: differential centrality-ensemble analysis based on gene expression profiles and protein-protein interaction network

**DOI:** 10.1101/2025.03.14.638654

**Authors:** Elnaz Pashaei, Sheng Liu, Kailing Li, Yong Zang, Lei Yang, Tim Lautenschlaeger, Jun Huang, Xin Lu, Jun Wan

## Abstract

Uncovering key genes that drive diseases and cancers is crucial for advancing understanding and developing targeted therapies. Traditional differential expression analysis often relies on arbitrary cutoffs, missing critical genes with subtle expression changes. Some methods incorporate protein-protein interactions (PPIs) but depend on prior disease knowledge. To address these challenges, we developed DiCE (Differential Centrality-Ensemble), a novel approach that combines differential expression with network centrality analysis, independent of prior disease annotations. DiCE identifies candidate genes, refines them with an information gain filter, and reconstructs a condition-specific weighted PPI network. Using centrality measures, DiCE ranks genes based on expression shifts and network influence. Validated on prostate cancer datasets, DiCE identified genes over-represented in key pathways and cancer fitness genes, significantly correlating with disease-free survival (DFS), despite DFS not being used in selection. DiCE offers a comprehensive, unbiased approach to identifying disease-associated genes, advancing biomarker discovery and therapeutic development.

**Graphical Abstract:** 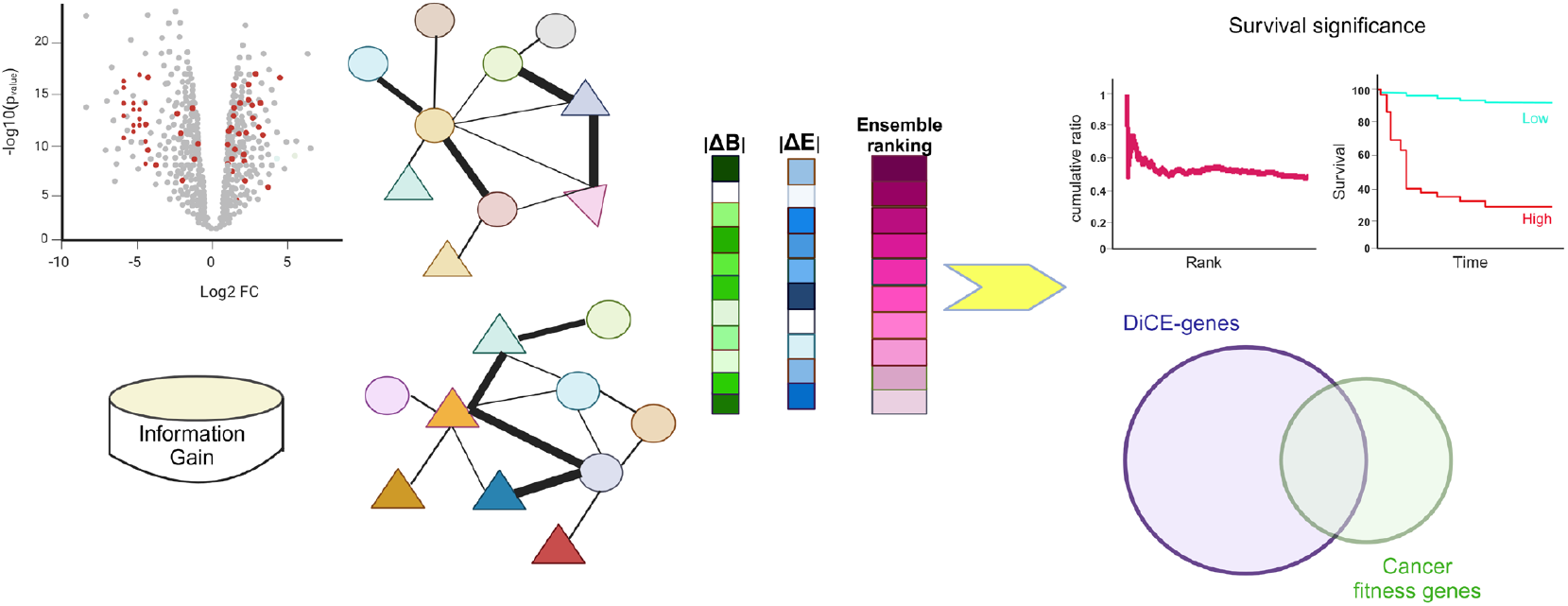

## Introduction

Differential expression analysis (DEA) is a pivotal step in pinpointing differentially expressed genes (DEGs) with expression levels systematically changed between two distinct conditions. Recently machine learning techniques have been increasingly applied to analyze genomic data for identifying DEGs. However, these methods predominantly rely on statistical models to analyze gene expression data and identify DEGs at the individual gene level. Especially, DEGs are often determined by arbitrarily chosen cutoffs, e.g., p-value or false discovery rate-adjusted p-value (q-value) below a specific value, e.g., 0.05, and/or expression fold change (FC) exceeding a predefined threshold (commonly |log_2_FC| > 1). While effective, such strict cutoffs risk overlooking biologically significant genes that might not exhibit substantial expression changes but play crucial roles in biological functions and pathways related to diseases or phenotypes. Alternative approaches, such as Weighted Gene Co-expression Network Analysis (WGCNA)[1], model-based gene clustering (MBCdeg)[2], and Bayesian model-based multi-tissue clustering algorithm (revamp)[3], categorize genes based on the similarity of expression patterns. However, focusing on individual gene levels without accounting for protein-protein interactions (PPIs) may fail to identify gene signatures with pivotal roles within the comprehensive gene regulatory network. Some methodologies such as PhenoGeneRanker[4], User-guided Knowledge-driven Integrative Network-based method (uKIN)[5], and Prioritization with a warped network (PWN)[6] employ disease-specific genes to navigate through the biological networks, e.g., PPIs, in order to prioritize genes associated with specific diseases. These approaches typically depend on a predefined set of disease-specific genes, often called “seed genes”, known to be associated with the disease under investigation[7]. If the information about seed genes is either absent or biased, the results may be inaccurate or incomplete.

To address these constraints and enhance the precision of gene prioritization to better identify key genes that are more contextually relevant with biological significance, we developed an innovative approach called <u>Di</u>fferential <u>C</u>entrality-<u>E</u>nsemble (DiCE). DiCE harnesses multiple elements, including DEA to construct a candidate gene pool, feature selection to choose the most informative genes filtered by the Information Gain (IG), followed by topological assessment of sample-specific weighted networks to capture context-specific interactions and utilization of ensemble ranking to combine multiple rankings, then prioritize genes identified by DiCE, namely DiCE-genes. This comprehensive framework enhances the robustness and interpretability of the gene selection process by integrating statistical significance with biological relevance.

The efficiency of DiCE was assessed with two prostate cancer (PCa) datasets, including PCa tumor versus normal samples and PCa metastatic tumors versus primary tumors. DiCE effectively identifies and prioritizes not only disease-candidate genes but also survival-correlated and fitness genes. These findings underscore DiCE’s potential to identify gene signatures linked to cancer mechanisms and prognostic significance, thus offering valuable insights into clinical applications in cancer management. Notably, DiCE operates independently of known disease-related genes, making it applicable even in contexts with limited prior knowledge.

## Materials and methods

### Workflow of DiCE

The DiCE algorithm mainly consists of six phases (Fig. 1). <u>Phase I</u>: Construction of a candidate gene pool by DEA with relaxed cutoffs, such as p < 0.05 with or without an FC threshold. <u>Phase II</u>: Selection of the top discriminative genes from the candidate pool obtained in Phase I using the Information Gain (IG) filter approach. <u>Phase III</u>: Modification of weighted PPIs network by calculating Pearson correlation coefficients (*c*.*c*.) of gene pairs in terms of gene expression for each specific phenotype or sample type. The value of (1 − |*c*· *c*· |) is assigned as the normalized distance between paired genes in accordance with the PPI network. <u>Phase IV</u>: Topological analysis of sample-specific weighted PPI networks from the perspectives of Betweenness and Eigenvector centrality, followed by differential analysis of each centrality measure for individual genes. <u>Phase V</u>: Ensemble ranking to produce a reliable and robust rating by integrating the ranks established in the previous stage. <u>Phase VI</u>: Enrichment analysis of biological pathways and functions for identified DiCE-genes.

**Figure 1.**
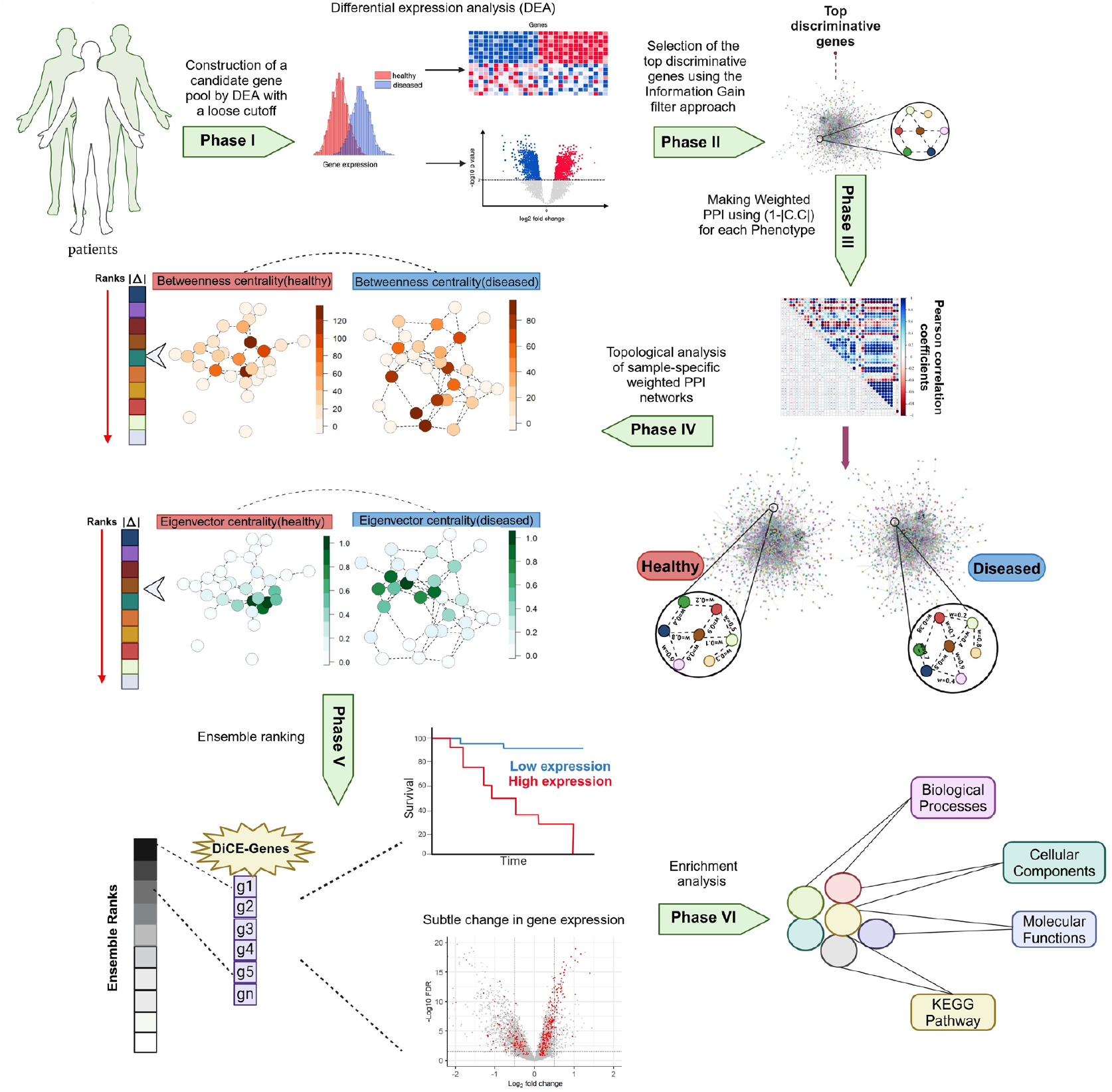
DiCE workflow.

### Phase I: Construction of gene candidate pool by DEA

DEA serves as the first step in the DiCE, generating a large candidate gene pool that forms the foundation for subsequent phases. In this study, DEA is performed using lmFit and eBayes functions in limma-voom[8] for the microarray dataset (GSE21032)[9] or edgeR package[10] in R for the RNA-seq data of the TCGA prostate adenocarcinoma (PRAD) cohort[11]. To ensure the inclusion of genes with subtle changes, loose cutoffs are applied, allowing small FCs and less statistical significance with higher p-values.

### Phase II: Selection of most discriminative genes using IG

The IG filter is a widely used statistical measure for gene selection[12]. It quantifies the amount of information a gene contributes to differentiating between conditions. Genes with higher IG values tend to be more informative and effective in distinguishing between groups.

Consider a dataset with *N* instances distributed across *k* classes. The ratio of samples in each class is denoted as 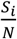, where *S*_*i*_ (*i* = 1, 2, · ·, *k*) represents the number of samples in the *i*^*th*^ class out of N total samples. The initial entropy *D* of the dataset, prior to any split, is calculated as follows:

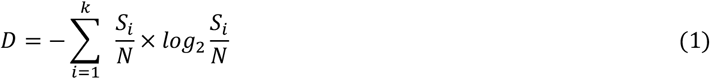

Given a gene *g* with different expression levels *V* = {*v*_1_, *v*_2_, …, *v*_*m*_}, where *v*_*i*_ denotes the distinct expression levels and *m* corresponds to the number of *v*_*i*_, the weighted entropy of gene *g* is calculated as follows:

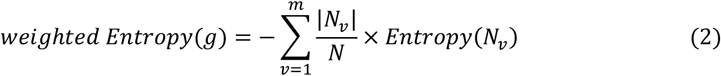

where *Nv* represents the partitioned subset of *N* samples with the size of |*N*_*v*_ | based on expression values of gene *g*. Finally, the IG value of gene *g* can be calculated as follows:

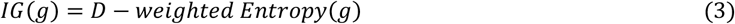

In fact, IG quantifies the reduction in entropy by measuring how well the expression levels of individual genes can partition samples into distinct classes. A higher IG indicates a greater ability to distinguish between classes, thereby reducing uncertainty (entropy) in classification.

The candidate gene pool refined by the IG filter keeps the most informative genes, characterized by their high discriminative powers. Specifically, genes are selected for the next analysis step if their IG values exceed the average IG value of all genes in the pool. This enhances interpretability, boosts classification performance, and optimizes computational efficiency, ultimately leading to a more effective analysis.

### Phase III: Modification of PPI network with gene expression correlations

The PPI networks provide a comprehensive view of the connections among genes. Genes are depicted as vertices linked by undirected edges that represent their interactions. Despite variations in biological systems being studied, the STRINGdb package[13] in R (v.11) for species 9606 with a confidence score threshold of 400 serves as the resource of PPIs.

To identify biologically relevant crucial genes among the top discriminative genes selected in Phase II, a systematic approach was employed according to their connectivity and interaction patterns. Given the Pearson correlation coefficients (*c*.*c*.) of gene expression for specific conditions or sample types between pairs of genes in Phase II, the value of (1 − |*c*· *c*· |) was used as a distance-like metric to characterize the degree of dissimilarity or lack of connectivity between nodes/genes to discover the shortest paths between network nodes. By generating condition/sample type-specific PPI networks, we can effectively capture the differentiation between various conditions or samples from the perspective of the strength of gene relationships within the networks.

### Phase IV: Topological analysis of sample-specific weighted PPI networks using betweenness and eigenvector centrality

Topological analysis examines various network metrics and properties to gain insights into its key structure and characteristics. For example, centrality metrics can pinpoint proteins that hold the most central or influential positions in the network and highlight proteins acting as bridges or are crucial for the flow of information.

In this study, two distinct centrality measures, betweenness, and eigenvector, were employed to assess node importance and significance within the condition/sample-specific weighted PPI networks. Betweenness centrality gauges the proportion of the number of shortest paths that traverse each node, providing an estimate of the frequency of the node lying on these paths between other nodes. The distances from or to the network’s vertices are used to calculate the shortest path. Higher betweenness centrality of one node indicates its greater impact on information flow within the network. It effectively identifies bottlenecks and evaluates the global importance of nodes. The robustness of betweenness centrality in the identification of target genes has been attested by many studies. Eigenvector centrality extends from the degree centrality by considering a node’s importance based on its connections to other important nodes. This metric accounts for both the number of connections and the importance and significance of nodes within the network by taking into consideration the degrees of neighboring nodes.

A node with numerous connections might have a relatively low eigenvector centrality if its connections link to nodes with low degrees. Similarly, a node could possess a high betweenness but a low eigenvector centrality, indicating that it is distant from the network’s influential nodes and bridges disparate parts within the network. Combining both eigenvector and betweenness centralities in network analysis provides complementary insights into node importance and influence and enhances our ability to identify and interpret key features within the network, thereby facilitating a deeper understanding of its intricate relationships, uncovering hidden patterns from gene regulatory network perspective, and elucidating the underlying mechanisms driving its behavior. By leveraging eigenvector and betweenness centralities, we are better equipped to discern critical genes, detect community structures, and grasp the flow of information or influence across the network.

### Phase V: DiCE-genes identified by gene prioritization using ensemble ranking based on absolute differences of each centrality measure

Considering the betweenness and eigenvector centralities of each gene within weighted PPI networks under distinct conditions or samples, such as normal and tumor samples or groups before and after treatment, we computed the absolute differences in each centrality measure between these various conditions/samples. Evaluating the absolute disparities in two centrality metrics enables us to spotlight genes that experienced notable shifts in their centralities, potentially indicating significantly altered roles or functional relevance within specific conditions or among distinct sample types. While both betweenness and eigenvector centralities were employed to evaluate condition/sample-specific weighted PPI networks, the absolute values of differences in either one of them yielded distinct rankings. This underscores the necessity of establishing an overall aggregated ranking of alternatives. Subsequently, the genes were ranked in descending order based on their absolute differences for each centrality measure. The genes with the greatest absolute differences were most likely important with pivotal roles associated with distinct groups/conditions, signifying their prominences and impacts within the context of condition or sample-specific weighted PPI networks, in addition to differential gene expression levels. Ensemble ranking, a technique in data analysis and decision-making, merges multiple individual rankings into a unified, aggregated ranking that enhances the overall accuracy, resilience, and reliability of the final gene ranking by harnessing the combined strengths of various ranking strategies. The DiCE-genes were identified after excluding those with centrality measures lower than the average value in any phenotype or condition.

### Phase VI: Biological function and pathway enrichment analysis

Enrichment analysis of biological functions and pathways was conducted to determine and interpret the comprehensive functional roles of DiCE-genes, providing more insights into the underlying molecular mechanisms responsible for the differences observed among distinct conditions/samples in the study. To identify Gene Ontology (GO) terms and KEGG pathways significantly enriched in the identified DiCE-genes, the interactive tool DAVID (Database for Annotation Visualization and Integrated Discovery, https://david.ncifcrf.gov/) was employed. Only GO terms and KEGG pathways with FDR less than 0.05 were considered significantly enriched.

### Survival analysis

The survival analysis was performed using the Kaplan-Meier (KM) log-rank test on all genes based on their gene expression levels, either high or low. Our study utilized disease-free survival (DFS) information of 500 PRAD patients in the “Prostate Adenocarcinoma (TCGA, Firehorse Legacy)” dataset[14] with corresponding clinicopathologic information available on cBioPortal.

## Results

### DiCE analysis on the TCGA PCa dataset

To perform the differential ensemble centrality analysis on the TCGA PCa dataset, we first obtained gene expression data from tumor and adjacent normal prostate samples using the TCGA platform (https://gdac.broadinstitute.org/)[11]. The dataset included 33 patients, each contributing tumor and paired normal tissue samples. Preprocessing steps involved removing noncoding genes (lncRNA and pseudogenes), resulting in an expression profile of 16,322 genes across tumor and normal samples. Fig. 2A summarizes the numbers of selected genes at each phase by the DiCE analysis applied to the TCGA PCa dataset. More details are provided below,

**Figure 2.**
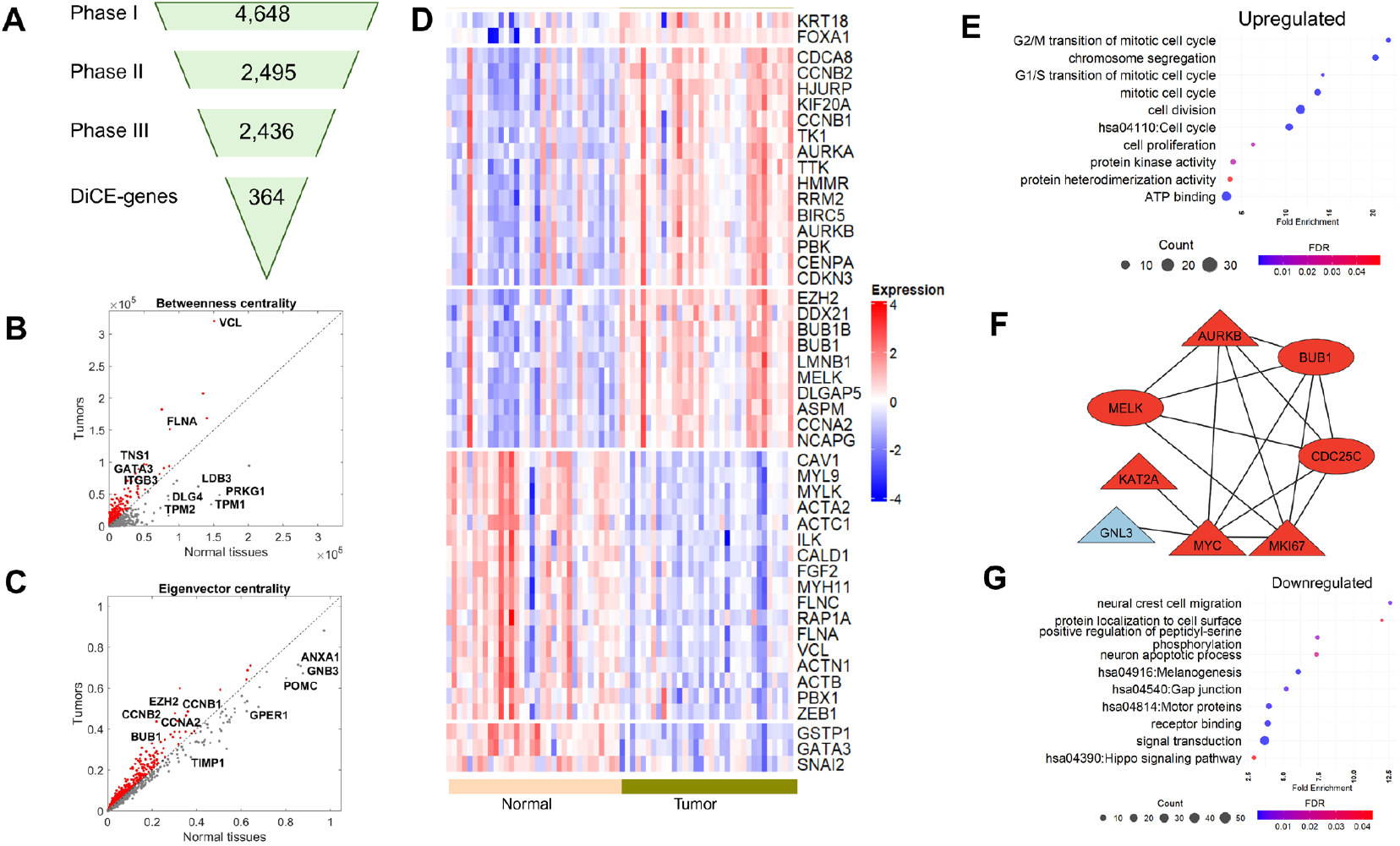
DiCE applied to PCa tumor vs. normal tissues. (**A)** Number of genes in each phase of the proposed approach. (**B)** and **(C)** Betweenness (**B**) and Eigenvector centrality (**C**) for each gene in normal tissues (x-axis) and PCa tumors (y-axis). **D**, Expression levels of DiCE-genes in normal tissues and tumors which were identified as a hub gene in previous studies. (**E)** Selected GO terms and KEGG pathway significantly enriched in upregulated DiCE-genes. (**F)** PPI network of upregulated DiCE-genes involved in cell proliferation. The node shapes represent changes in gene betweenness between tumor and normal samples (triangle for an increase and oval for a decrease), while the red and blue colors indicate a positive or negative shift in eigenvector centrality, respectively. (**G)** GO/KEGG pathways notably over-represented in downregulated DiCE-genes.

**Phase I** constructed the pool of candidate genes. DEA was performed using the edgeR[10] for paired comparison between PCa tumors and adjacent normal samples to select candidate genes based on loose cutoffs: FDR < 0.05 and |log_2_FC| >0.5, resulting in 4,648 candidate genes in the pool (Fig. 2A). Z-score normalization was then applied to the expression data of selected candidate genes for downstream analysis.

In **Phase II**, the IG filter was applied to all candidate genes to identify the most discriminative genes from the candidate pool. Each gene was assigned a weight based on its entropy. A total of 2,495 genes with weights higher than the mean weight were identified as the most discriminative genes for further analysis (Fig. 2A).

In **Phase III**, the PPI network of the most discriminative genes based on the STRING database[13] was adjusted based on gene expression correlations across different conditions or samples. Correlations were treated as a distance-like metric, calculated as (1 − |*c*· *c*· |), in the PPI network. After excluding 59 genes that lacked connections, the resulting sample-specific weighted PPI networks included 2,436 genes (Fig. 2A) and 19,472 interactions.

**Phase IV** included a topological analysis of the sample-specific weighted PPI networks using two centrality measures: betweenness and eigenvector. These parameters were calculated for each gene (node) within the network for both tumor and normal samples.

An increase in the betweenness centrality of a node/gene in tumor samples indicates that the gene lies on shorter paths connecting other genes, suggesting its role as a bridge in tumor-related PPIs (Fig. 2B), for instance, VCL, FLNA, TNS1, GATA3, and ITGB3 which may act as key connectors in tumor samples. Conversely, decreased betweenness centralities observed for genes like LDB3, PRKG1, TPM1, TPM2, and DLG4 suggest reduced involvement in shortest paths or a diminished role in connecting to other genes (Fig. 2B).

Eigenvector centrality measures the relative influence of a node/gene within the network (Fig. 2C). An increase in eigenvector centrality indicates connections to highly influential genes, enhancing the gene’s significance in the network as EZH2, CCNB1, CCBB2, CCNA2, and BUB1 in PCa tumors. On the other hand, a decrease in eigenvector centrality indicates that a gene is connected to less influential or weakening genes, possibly resulting in lower relative influence scores or reduced significance in the network, such as ANXA1, GNB3, POMC, GPER1, and TIMP1 in tumor samples (Fig. 2C).

In **Phase V**, ensemble ranking was used to prioritize genes based on the absolute differences of each centrality measure from Phase IV. Genes were excluded if their centrality measures in any sample groups/types/conditions were below the average value for the corresponding group. This process identified 364 DiCE-genes, comprising 138 up-regulated and 226 down-regulated genes in PCa tumors. These DiCE-genes demonstrated notable alterations of these two centrality scores, along with gene expression changes between tumor and normal samples, highlighting their crucial role in shaping the network structure and influencing other genes in response to condition changes. Among them, 47 DiCE-genes were recognized as hub genes (Fig. 2D) in multiple PCa studies[15-24].

We performed functional enrichment analysis on up- and down-regulated DiCE-genes in terms of GO and KEGG pathways. The 138 up-regulated genes were enriched in processes such as cell proliferation, cell cycle, cell division, chromosome segregation, protein kinase activity, ATP binding, protein heterodimerization activity, and multiple mitotic-related biological processes, e.g., mitotic cell cycle (Fig. 2E). Eight of these up-regulated DiCE-genes, KAT2A, MELK, MYC, MKI67, CDC25C, BUB1, AURKB, and GNL3, are associated with cell proliferation (Fig. 2F). Among them, five genes showed higher betweenness (triangle) in PCa tumor and almost all of them exhibited increased eigenvector centrality (red color). These findings underscore their roles as central hubs orchestrating broader transcriptional, signaling, or regulatory changes.

In contrast, 226 down-regulated genes were associated with multiple KEGG pathways such as Melanogenesis, Gap junction, Hippo signaling pathways, and biological processes, e.g., signal transduction, receptor binding, neural crest cell migration, protein location to cell surface, neuron apoptotic process, etc. (Fig. 2G). The Hippo pathway is crucial in PCa and modulates AR signaling via YAP, impacting AR+ PCa growth[25]. Additionally, 12 out of 15 downregulated motor proteins exhibited reduced betweenness centrality in PCa tumors, including TPM1 and TMP2 (Fig. 2B), both recognized as tumor suppressors, further indicating a disruption of tumor-suppressor circuits during PCa progression and highlighting potential therapeutic targets.

### DiCE analysis on PCa metastatic data

We applied DiCE to the GSE21032[9] dataset, comprising 131 primary and 19 metastatic PCa tumors. DEA by limma identified 7,757 candidate genes with a loose cutoff of p-value < 0.05 (Fig. 3A). A subset of 3,403 genes was identified as the most discriminative in Phase II using the IG filter, followed by reconstruction of the PPI network (Phase III) keeping 3,186 genes with modified node distances using (1 - |*c*.*c*.|) based on gene expression profiles in PCa metastatic and primary tumors, respectively. Topological differential analysis in Phase IV further narrowed the list to 347 DiCE-genes in Phase V (Fig. 3A), encompassing 47 genes identified through a meta-analysis of five studies[26] and hub genes (Fig. 3B).

**Figure 3.**
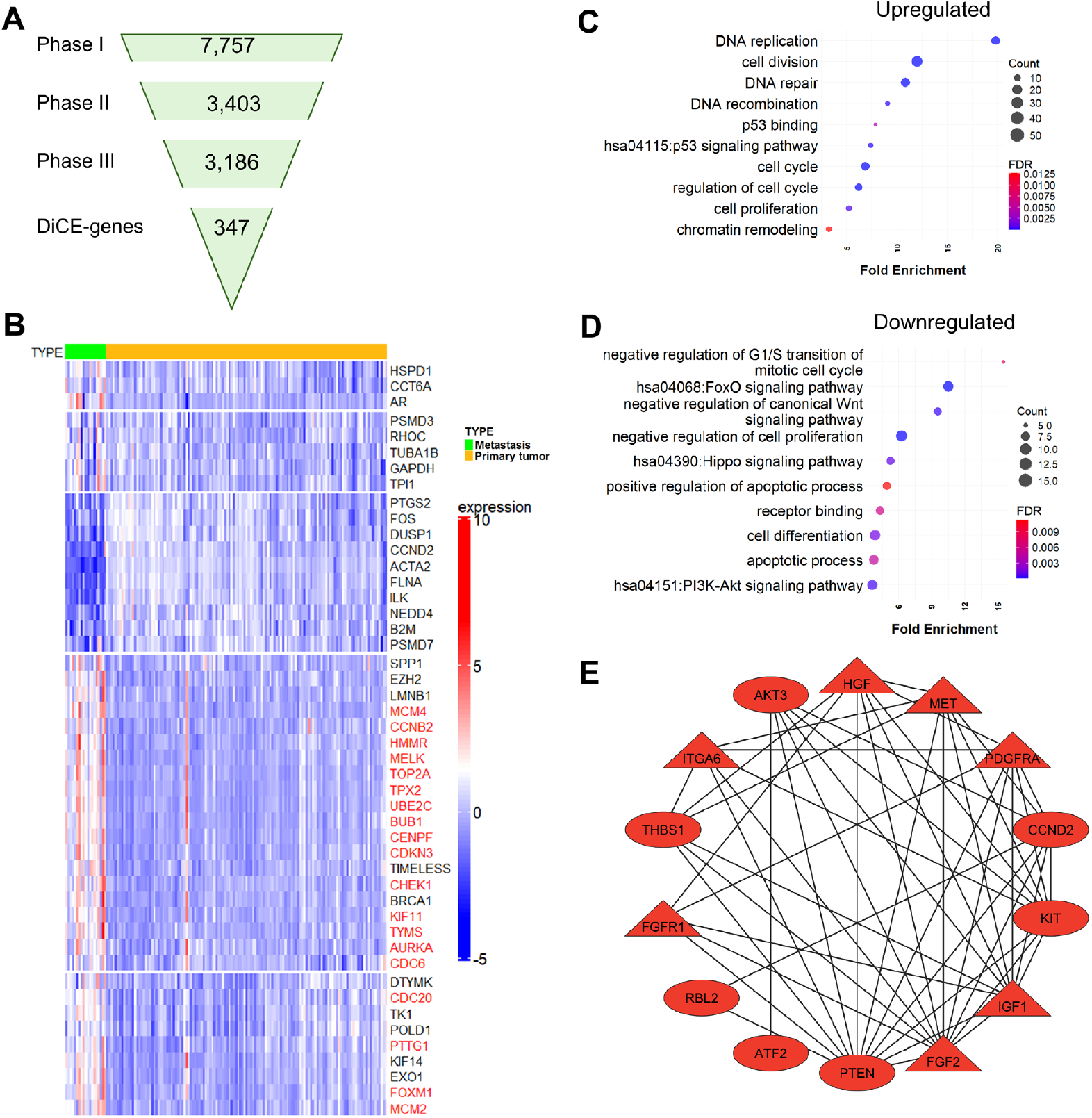
DiCE applied to PCa metastasis vs. primary tumors. **(A)** Number of genes in each phase of DiCE. (**B)** Expression of 47 selected DiCE-genes associated with PCa metastasis identified through a meta-analysis of five studies. Genes highlighted in red denote hub genes. (**C)** and **(D)** Enriched GO terms and KEGG pathway among upregulated (**C**) and downregulated (**D**) DiCE-genes. (**E)** PPI network for downregulated DiCE-genes in the PI3K-Akt signaling pathway. (**G)** Overlap of two sets of DiCE-genes based on PCa tumor vs normal tissues and metastasis vs primary tumors. (**F)** Ratios of DFS-related DiCE-genes in each segment of (**G**).

238 upregulated DiCE-genes in PCa metastasis were significantly enriched in cell cycle, cell division, cell proliferation, DNA repair/replication/recombination, and chromatin remodeling (Fig. 3C). Strikingly, several genes involved in p53 binding and p53 signaling pathway, including GTSE1, RRM2, CCNB1, and CHEK1, were upregulated in metastatic PCa. GTSE1, which is highly expressed in various cancers, including PCa, and linked to poor prognoses[27], promotes PCa cell proliferation by activating the SP1/FOXM1 signaling pathway[28]. Similarly, RRM2 overexpression was associated with aggressive PCa and unfavorable outcomes[29]. Despite their elevated expression, both GTSE1 and RRM2 exhibited reduced betweenness centrality in PCa metastasis compared to primary tumors. Moreover, the combined overexpression of RRM2 and CCNB1 has been implicated in driving PCa progression via the cell cycle and the p53 signaling pathway[29]. While CCNB1 and CHEK1 showed decreased eigenvector centrality in metastasis, CHEK1 plays a key role in the DNA damage response[30] and contributes to tumor progression and therapy resistance in PCa, particularly in metastatic castration-resistant prostate cancer (mCRPC)[31].

The 109 downregulated DiCE-genes are involved in a wide range of biological processes (Fig. 3D), e.g., apoptotic process, cell differentiation, receptor binding, negative regulation of cell proliferation, negative regulation of canonical Wnt signaling pathway, besides Hippo, FoxO, and PI3K-Akt signaling pathways. 14 of these DiCE-genes, PDGFRA, ATF2, HGF, PTEN, IGF1, FGF2, THBS1, RBL2, CCND2, KIT, AKT3, ITGA6, MET, and FGFR1, are linked to the PI3K-Akt pathway, a key oncogenic driver of migration, proliferation, and drug resistance, frequently dysregulated in metastatic PCa[32]. Among them, PTEN is a well-established tumor suppressor in PCa, acting as a major inhibitor of growth signaling and interacting with the MAPK pathway through PPIs[33]. Oncogenes, such as FGF2, FGFR1, HGF, IGF1, MET, and PDGFRA, showed higher betweenness (triangles in Fig. 3E) in metastatic PCa, despite their reduced gene expression. Additionally, all 14 DiCE-genes displayed increased eigenvector centrality (red color in Fig. 3E), emphasizing their enhanced influence within the network during PCa metastasis.

### DiCE-genes are more likely correlated with patient survival outcomes

Next, survival analysis was conducted on DiCE-genes using the Cox proportional hazards model. Among 364 DiCE-genes identified upon the comparison between PCa tumors and normal samples, 134 (36.8%) presented significant associations (p < 0.05) between their expression levels and patients’ DFS, even though DFS information was not included during DiCE-gene selection. Cumulative ratios of DiCE-genes associated with DFS outcomes, along with their final DiCE rankings, revealed that lower gene ensemble ranks led to a decline in the proportion of survival-correlated genes (Fig. 4A). These suggest that top-ranked DiCE-genes tend to be more correlated with clinical outcomes, such as CCNB2 (rank #1), EZH2 (rank #2), CALM1 (rank #4) (Fig. 4B). Cyclin B2 (CCNB2), a key regulator of the cell cycle, has been reported to be upregulated in human cancers. Its increased expression is associated with poor survival outcomes, suggesting its potential as a novel prognostic maker. EZH2 plays a multifaceted role in PCa progression. Many studies have demonstrated that the polycomb repressive complex 2 (PRC2)-dependent transcription repression by EZH2 suppresses interferon γ-signaling and promotes PCa cell invasion, cancer stem cell features, and angiogenesis, among others. The aberrant expression of CALM1 was detected in several cohorts of PCa patients.

**Figure 4.**
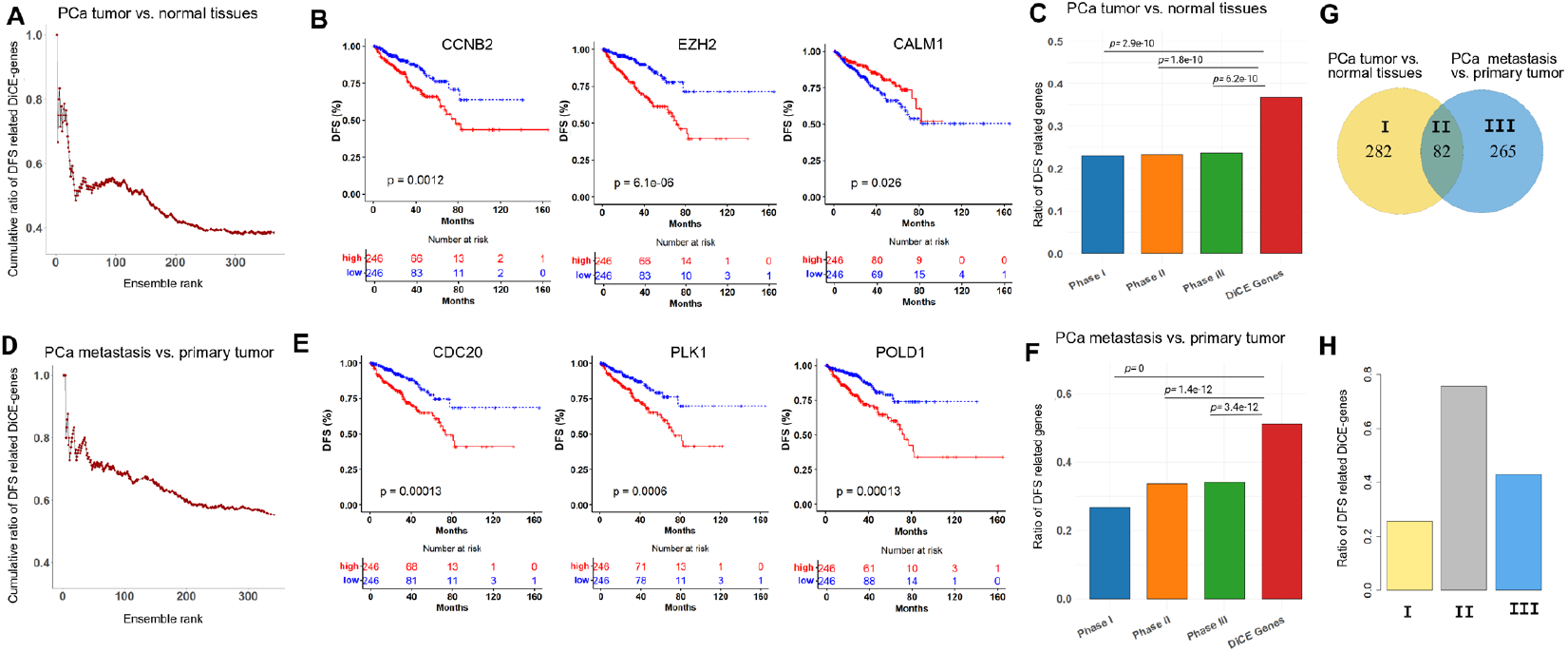
Association between key genes and patients’ survival outcomes. (**A)** Cumulative ratios of DiCE-genes identified between PCa tumor and normal tissues which were significantly associated with DFS along with their ensemble rankings. (**B)** Kaplan-Meier (KM) plots of top-ranked DiCE-genes with significant survival outcomes. **C**, Ratios of DFS-related genes in each phase of DiCE. **(D)** Cumulative ratios of DFS-related DiCE-genes for PCa metastasis vs. primary tumors along with their rankings. (**E)** KM plots of selected top-ranked DiCE-genes. **F**, Ratio of DSF-related genes in each DiCE phase. The statistical significance of the ratio differences in (**C**) and (**F**) was evaluated based on hypergeometric distribution.

For comparison, ratios of genes linked to patient outcomes at each phase of the DiCE were calculated (Fig. 4C). Only 22.4% of 4,648 candidate genes after Phase I (DEA) showed survival significance, along with 22.9% and 23.3% of genes from Phase II and III of DiCE, respectively. All of them were notably lower than the ratio of DiCE-genes according to the hypergeometric test.

A similar trend was observed for the DiCE-genes distinguishing metastasis and primary PCa. 51.0% of 347 DiCE-genes were significantly associated with DFS outcomes (Fig. 4D), including top ranked DiCE-genes, such as CDC20 (rank #1), PLK1 (rank #3), and POLD1 (rank #14) (Fig. 4E). CDC20 and PLK1 are important biomarkers and potential therapeutic targets for metastatic PCa. The diverse expression profiles of POLD1 (DNA polymerase delta 1 catalytic subunit gene) in various tumor types indicate its potential significance across different cancers as reported in previous studies, even though its specific role in PCa remains less defined. However, in our study, POLD1 explicitly exhibited a favorable rank and a highly significant survival p-value, indicating its potential as a prognostic marker or therapeutic target in PCa. Again, the survival significance ratio of DiCE-genes in Phase V was considerably higher than in any prior phases (Fig. 4F).

Among the 347 DiCE-genes distinguishing PCs metastatic from primary tumors, 82 were also recognized as DiCE-genes in the comparison between PCa tumors and normal samples (Fig. 4G). The significant overlap highlights the dual roles of genes such as TOP2A, CCNB2, BUB1, TPX2, and EZH2 in both tumor initiation and tumor progression, reinforcing their potential as key biomarkers for PCa. This consistent identification across different comparisons strengthens their relevance for further experimental validation and therapeutic exploration. Importantly, 62 out of these 82 shared DiCE-genes (75.6%) were significantly associated with disease-free survival (Fig. 4H), emphasizing their strong prognostic potential and roles in understanding PCa mechanisms. In comparison, lower proportions of DiCE-genes linked to survival significance were observed exclusively in PCa tumor vs normal (72 out of 282, 25.5%) or PCa metastasis vs primary tumor (114 out of 265, 43.0%), suggesting distinct genetic landscapes underpinning various PCa subtypes (Fig. 4H).

### DiCE-genes not discovered by canonical DEA with strict cutoffs

Traditional DEG analysis typically focuses on genes that exhibit the most significant changes in expression levels. Genes that do not meet certain statistical criteria may be overlooked. However, cancer is a complex disease, and genes involved in its progression may have subtle or context-dependent roles that are not reflected in simple changes in expression levels. For example, 163 out of 364 DiCE-genes determined by the comparison between PCa tumor and normal samples can’t pass the common cutoff for conventional DEA, e.g., |log_2_FC| > 1 (Fig. 5A), comprising 80 upregulated and 83 downregulated genes.

**Figure 5.**
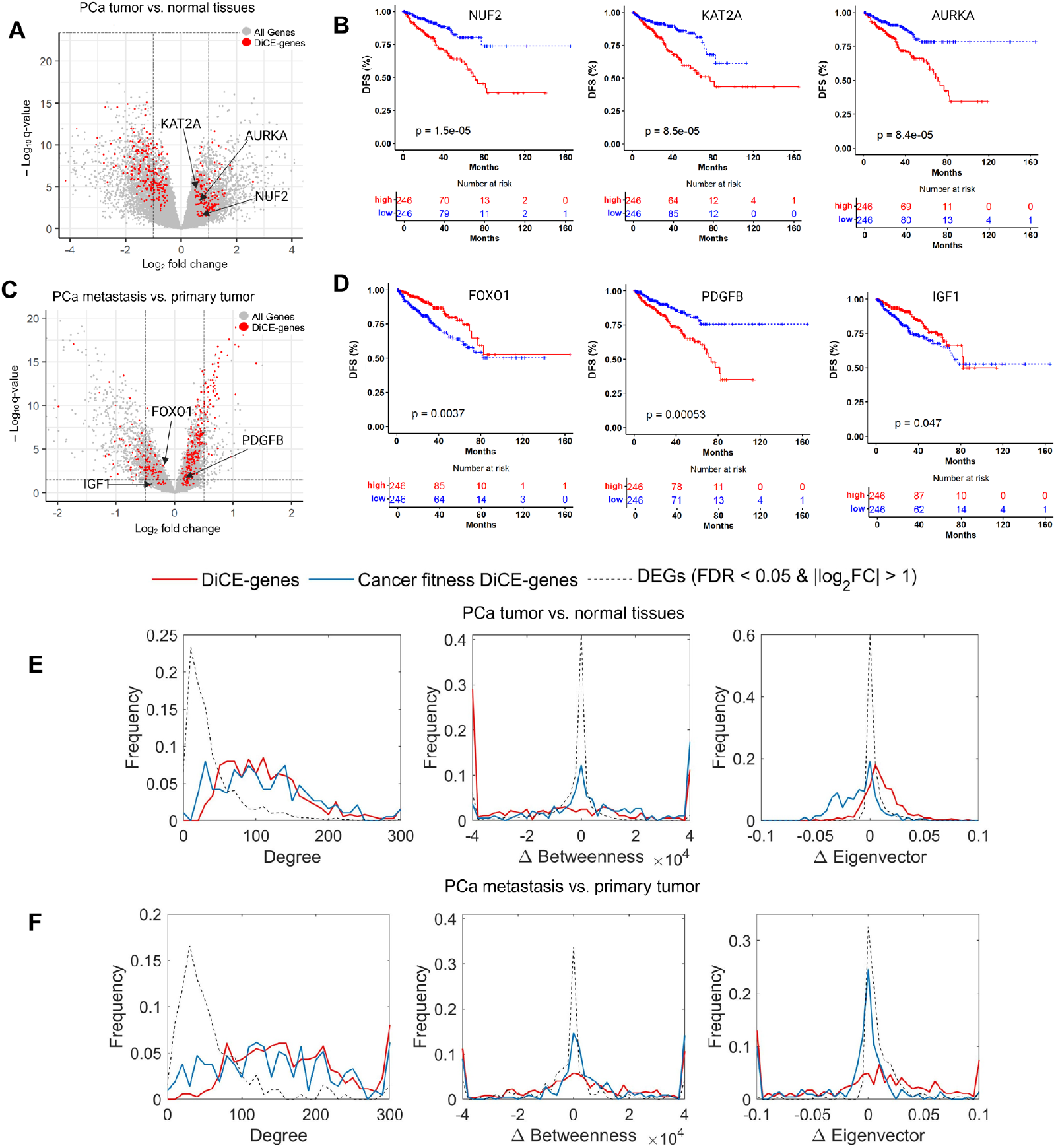
DiCE-genes not identified by traditional DEA. **(A)** Volcano plot highlighting three DiCE-genes with minimal expression changes between PCa tumors and normal tissues. (**B)** KM plots of three genes selected. (**C)** Volcano plot for PCa metastasis vs. primary tumors, highlighted by three genes with subtle gene expression changes. (**D)** KM plots for three DiCE-genes in (**C**). (**E)** and (**F)** Distribution of degree, Δbetweeness, and Δeigenvector centrality for DiCE-genes (red line), cancer fitness DiCE-genes (blue line), and DEGs (dotted line), in the study of PCa tumor/normal tissues (**E**) and PCa metastasis/primary tumors (**F**), respectively.

These DiCE-genes were enriched in dysregulation of pathways common in PCa and contribute to aggressiveness and treatment resistance, suggesting their important roles in tumor progression and potential as therapeutic targets in PCa. It’s important to note that the top-ranked DiCE-genes don’t necessarily exhibit the highest statistical significance or the largest FCs in expression. Some key genes might have been excluded by the stringent cutoff criteria typically used in standard analyses. For example, NUF2, KAT2A, and AURKA had relatively modest log_2_FC values of 0.69, 0.52, and 0.70, respectively. KAT2A (lysine acetyltransferase 2A) plays a crucial role in PCa by modulating androgen receptor (AR) activity through acetylation.

Its overexpression correlates with resistance to hormonal therapy, contributing to disease progression and poor clinical outcomes[34]. Targeting KAT2A holds a promising strategy to overcome treatment resistance in CRPC[34]. The dysregulation of AURKA (Aurora kinase A) plays a crucial role in the tumorigenesis and development of various cancers, including PCa[35]. The observed overexpression of AURKA in CRPC samples, especially in cases with amplification of the AR gene or elevated AR expression levels, highlights its potential significance as a therapeutic target in AR-positive CRPC[36,37]. NUF2, a component of the NDC80 kinetochore complex, is essential for proper chromosome segregation during the mitotic process[38]. While its direct involvement in PCa remains underexplored, studies in other cancers suggest its role in genomic instability and tumor progression, including breast cancer, lung cancer, and clear cell renal cell carcinoma[39-41]. Aberrant NUF2 expressions or activities may contribute to PCa aggressiveness, metastasis, or therapeutic resistance, warranting further investigation. Moreover, these genes were significantly associated with survival outcomes (Fig. 5B). Notably, 49 out of 134 survival-related DiCE-genes exhibited only minimal FCs in expression (|log_2_FC| < 1), a detail that might be missed by conventional DEG methods. Importantly, the identification of these genes by DiCE underscored their potential as PCa biomarkers, highlighting the effectiveness of our approach in uncovering critical genes with subtle yet meaningful contributions to disease progression that may be overlooked by traditional approaches.

The comparison between metastatic and primary PCa tumors also revealed that 238 out of 347 DiCE-genes exhibited log_2_FC amplitudes smaller than 0.5, with some showing |log_2_FC| < 0.2 (Fig. 5C), including top-ranked genes such as CDC20 (rank #1 with log_2_FC = 0.45, Fig. 4E), and survival-associated genes like PDGFB and FOXO1 (Fig. 5D). We also observed 23 DiCE-genes with *q*-values > 0.05 in DEA, likely due to expression variations or tumor heterogeneity. Among them, IGF1 was strongly associated with patient survival outcomes (Fig. 5D), consistent with previous reports supporting its important role in PCa[42]. More examples include CDK13 (*q* = 0.076), CSNK2A2 (*q* = 0.093), DECR1 (*q* = 0.072), PARP1 (*q* = 0.069), and PRMT5 (*q* = 0.13).

Studies have shown that CDK13 promotes lipid deposition and PCa progression by stimulating NSUN5-mediated m5C modification of ACC1 mRNA[43]. CSNK2A2 has been reported as a marker of PCa progression and prognosis[44], while DECR1 acts as an androgen-repressed survival factor that regulates PUFA oxidation to protect prostate tumor cells from ferroptosis[45]. PARP1 is a well-established therapeutic target in cancers including PCa, especially in patients harboring BRCA mutations[46]. PRMT5 overexpression has been closely linked to AR expression at both the protein and mRNA levels. Knockdown of PRMT5 specifically inhibited the growth of PCa cells in an AR-dependent manner and suppressed AR transcription[47,48]. These emphasize the limitations of conventional DEG methods in identifying important genes solely based on DEA.

As previously identified, 36.8% of DiCE-genes distinguishing PCa tumors from normal tissues were significantly associated with survival outcomes, a considerably higher proportion than observed for DEGs (23.1%). Among the 134 survival-associated DiCE-genes, 76 exhibited a hazard ratio (HR) below 1, indicating that lower expression levels were linked to improved survival. Notably, 71 out of these 76 DiCE-genes were upregulated in PCa tumors compared to adjacent normal tissues. Conversely, 51 of the 58 DiCE-genes with HR above 1 were downregulated in PCa tumors. Even more strikingly, 154 out of 155 DiCE-genes with HR < 1 were upregulated in PCa metastasis compared to primary tumors, while all 22 DiCE-genes with HR > 1 were downregulated in metastatic tumors.

Cancer fitness genes contribute to the survival, proliferation, and adaptability of cancer cells. Unlike classical oncogenes and tumor suppressor genes, they may not initiate tumor formation but are essential for sustaining tumor growth, evading immune responses, and developing therapy resistance. Large-scale functional screens, such as CRISPR and RNAi-based knockout studies, have recently identified cancer fitness genes[49-53]. A total of 1,927 fitness genes were identified across all three PCa cell lines, 22RV1, DU-145, and LNCaP-Clone-FGC[50]. Among them, 575 and 577 were present in 16,322 and 19,302 genes detected in two datasets used in our study, accounting for 3.5% and 3.0%, respectively. Notably, these fitness genes were significantly over-represented in the identified DiCE-genes, comprising 6.3% (*p* = 2.6×10^−3^) and 15.9% (*p* = 1.5×10^−24^), whereas only 0.6% and 1.9% of traditional DEGs were classified as cancer fitness genes. These findings indicate the functional significance of the DiCE-genes in cancer development and progression, offering valuable insights into potential therapeutic targets for precision medicine and cancer treatment.

Furthermore, DiCE-genes, including those classified as cancer fitness genes, demonstrated greater connectivity within the PPI network compared to DEGs identified by conventional DEA, even when exhibiting only modest or minimal changes in gene expression (Figs 5E&F). They also displayed more pronounced alterations in the network structure, as reflected by two centrality measures such as betweenness and eigenvector centrality (Figs 5E&F).

All these findings highlight the advantage of our proposed DiCE approach in pinpointing key genes for distinguishing different conditions or phenotypes, such as tumor versus normal, or metastasis versus primary tumors, while also revealing their clinical significance in patient survival outcomes. Certain DiCE-genes that might be overlooked by traditional DEA due to typical cutoffs could still play important roles in cancer progression through diverse mechanisms. Their importance lies in their associations with, or co-regulation by, other genes within the PPI network, that DiCE can capture through alterations of centrality measures. Hence, DiCE provides a more comprehensive understanding of disease mechanisms and helps uncover novel therapeutic targets.

## Discussion

In this study, we introduced DiCE, a novel computational framework designed for gene prioritization and biomarker discovery without requiring prior knowledge of disease-causing genes. DiCE leverages multiple analytical techniques, including feature selection, topology analysis of sample-specific weighted PPI networks, centrality measures, and ensemble ranking. As a comprehensive framework, DiCE performs well in analyzing gene expression changes within the context of biological networks. Its versatility enables application across a wide range of diseases and biological processes, allowing researchers to utilize the approach with various datasets and experimental designs to enhance gene prioritization and uncover biologically critical genes, termed DiCE-genes, for specific conditions or phenotypes.

When applied to two PCa datasets, DiCE demonstrated its robustness, reliability, and effectiveness in identifying and prioritizing two sets of DiCE-genes, 364 genes distinguishing PCa tumor from adjacent normal tissues and 347 genes differentiating PCa metastatic from primary tumors, respectively. A total of 80 DiCE-genes were common across both datasets. Among the 347 DiCE-genes identified in metastatic PCa, 47 overlapped with genes from a meta-analysis of five studies, indicating their potential role in PCa metastasis, particularly in bone, lymph node, and liver metastasis. Notably, 19 of these DiCE-genes were highlighted as hub genes in the same meta-analysis, reinforcing the effectiveness and reliability of the DiCE approach.

DiCE successfully identified well-known PCa biomarkers, such as AR, TOP2A, and TPX2, while also capturing a significant number of potential marker genes with smaller FC amplitudes and/or lower statistical significance, such as KAT2A, AURKA, NUF2, CDC20, PDGFB, IGF1, and FOXO1, which could be missed by conventional DEG analysis methods.

Importantly, 36.8% and 51.0% of the two sets of DiCE-genes, including those with subtle expression changes, showed strong correlations with DFS outcomes, particularly among 82 shared DiCE-genes for both studies (75.6%). These results outperformed those achieved by conventional DEG analysis methods. Additionally, 91.8% and 99.4% of these DFS-linked DiCE-genes exhibited a strong concordance between their expression changes and HRs. DiCE-genes with HR < 1 were predominantly upregulated in PCa tumors compared to adjacent normal tissues, or in PCa metastasis compared to primary tumors, whereas genes with HR > 1 were downregulated. Cancer fitness genes, which promote tumor cell survival and proliferation by alleviating cellular stress, are promising targets for cancer therapies. We found that 6.3% and 15.9% of DiCE-genes were identified as fitness genes across all three PCa cell lines, far exceeding 0.6% and 1.9% observed in traditional DEGs. This further underscores the significance of DiCE-genes in shaping survival outcomes of PCa patients and highlights their potential as diagnostic and prognostic biomarkers, as well as therapeutic targets. Additionally, the results demonstrate DiCE’s precision in identifying biologically relevant genes, effectively linked to cancer fitness genes and patient survival outcomes compared to traditional DEG analysis, which often yields broader, less focused targets.

In conclusion, our results indicate the significance of adopting a comprehensive and multi-dimensional approach to differential expression analysis. Relying solely on traditional DEG analysis risks overlooking important genes and underlying mechanisms. By incorporating differential analysis of PPI networks, DiCE provides novel insights into disease biology, offering an advanced strategy for identifying clinically relevant genes. The approach prioritizes genes based on centrality measure changes, rescuing important marker genes that might be missed by conventional methods. DiCE’s versatility and broad applicability make it a powerful tool for studying diverse diseases and biological contexts. However, its success depends on factors such as the quality and completeness of PPI networks and the availability of functional gene data. Additionally, challenges related to multimodal data and heterogeneous information may arise. Incorporating tailored strategies for different data types and integrating them into the analytical pipeline can enhance accuracy and facilitate the exploration of biological mechanisms.

## Data availability

All data are available from the authors upon request.

## References

1. Mohammad, T., Singh, P., Jairajpuri, D.S. et al. Differential Gene Expression and Weighted Correlation Network Dynamics in High-Throughput Datasets of Prostate Cancer. Front Oncol. 2022; 12: 881246. 10.3389/fonc.2022.881246.

2. Osabe, T., Shimizu, K. and Kadota, K. Differential expression analysis using a model-based gene clustering algorithm for RNA-seq data. BMC Bioinformatics. 2021; 22: 511. 10.1186/s12859-021-04438-4.

3. Erola, P., Bjorkegren, J.L.M. and Michoel, T. Model-based clustering of multi-tissue gene expression data. Bioinformatics. 2020; 36: 1807–1813. 10.1093/bioinformatics/btz805.

4. Dursun, C., Kwitek, A.E. and Bozdag, S. PhenoGeneRanker: Gene and Phenotype Prioritization Using Multiplex Heterogeneous Networks. IEEE/ACM Trans Comput Biol Bioinform. 2022; 19: 2950–2962. 10.1109/TCBB.2021.3098278.

5. Hristov, B.H., Chazelle, B. and Singh, M. uKIN Combines New and Prior Information with Guided Network Propagation to Accurately Identify Disease Genes. Cell Syst. 2020; 10: 470–479 e473. 10.1016/j.cels.2020.05.008.

6. Han, S., Hong, J., Yun, S.J. et al. PWN: enhanced random walk on a warped network for disease target prioritization. BMC Bioinformatics. 2023; 24: 105. 10.1186/s12859-023-05227-x.

7. Liu, S., Nam, H.S., Zeng, Z. et al. CDHu40: a novel marker gene set of neuroendocrine prostate cancer. Brief Bioinform. 2024; 25. 10.1093/bib/bbae471.

8. Ritchie, M.E., Phipson, B., Wu, D. et al. limma powers differential expression analyses for RNA-sequencing and microarray studies. Nucleic Acids Res. 2015; 43: e47. 10.1093/nar/gkv007.

9. Taylor, B.S., Schultz, N., Hieronymus, H. et al. Integrative genomic profiling of human prostate cancer. Cancer Cell. 2010; 18: 11–22. 10.1016/j.ccr.2010.05.026.

10. Robinson, M.D., McCarthy, D.J. and Smyth, G.K. edgeR: a Bioconductor package for differential expression analysis of digital gene expression data. Bioinformatics. 2010; 26: 139–140. 10.1093/bioinformatics/btp616.

11. Cancer Genome Atlas Research, N., Weinstein, J.N., Collisson, E.A. et al. The Cancer Genome Atlas Pan-Cancer analysis project. Nat Genet. 2013; 45: 1113–1120. 10.1038/ng.2764.

12. Gao, L., Ye, M., Lu, X. et al. Hybrid Method Based on Information Gain and Support Vector Machine for Gene Selection in Cancer Classification. Genomics Proteomics Bioinformatics. 2017; 15: 389–395. 10.1016/j.gpb.2017.08.002.

13. Szklarczyk, D., Gable, A.L., Nastou, K.C. et al. The STRING database in 2021: customizable protein-protein networks, and functional characterization of user-uploaded gene/measurement sets. Nucleic Acids Res. 2021; 49: D605–D612. 10.1093/nar/gkaa1074.

14. Gao, J., Aksoy, B.A., Dogrusoz, U. et al. Integrative analysis of complex cancer genomics and clinical profiles using the cBioPortal. Sci Signal. 2013; 6: pl1. 10.1126/scisignal.2004088.

15. Song, Z.Y., Chao, F., Zhuo, Z. et al. Identification of hub genes in prostate cancer using robust rank aggregation and weighted gene co-expression network analysis. Aging (Albany NY). 2019; 11: 4736–4756. 10.18632/aging.102087.

16. Song, Z., Huang, Y., Zhao, Y. et al. The Identification of Potential Biomarkers and Biological Pathways in Prostate Cancer. J Cancer. 2019; 10: 1398–1408. 10.7150/jca.29571.

17. Khosravi, P., Gazestani, V.H., Akbarzadeh, M. et al. Comparative Analysis of Prostate Cancer Gene Regulatory Networks via Hub Type Variation. Avicenna J Med Biotechnol. 2015; 7: 8–15.

18. Li, S., Hou, J. and Xu, W. Screening and identification of key biomarkers in prostate cancer using bioinformatics. Mol Med Rep. 2020; 21: 311–319. 10.3892/mmr.2019.10799.

19. Singh, A.N. and Sharma, N. Identification of key pathways and genes with aberrant methylation in prostate cancer using bioinformatics analysis. Onco Targets Ther. 2017; 10: 4925–4933. 10.2147/OTT.S144725.

20. Liu, S., Wang, W., Zhao, Y. et al. Identification of Potential Key Genes for Pathogenesis and Prognosis in Prostate Cancer by Integrated Analysis of Gene Expression Profiles and the Cancer Genome Atlas. Front Oncol. 2020; 10: 809. 10.3389/fonc.2020.00809.

21. Zhu, H., Lin, Q., Gao, X. et al. Identification of the hub genes associated with prostate cancer tumorigenesis. Front Oncol. 2023; 13: 1168772. 10.3389/fonc.2023.1168772.

22. Gu, P., Yang, D., Zhu, J. et al. Bioinformatics analysis identified hub genes in prostate cancer tumorigenesis and metastasis. Math Biosci Eng. 2021; 18: 3180–3196. 10.3934/mbe.2021158.

23. Mukherjee, S. and Sudandiradoss, C. Transcriptomic analysis of castration, chemo-resistant and metastatic prostate cancer elucidates complex genetic crosstalk leading to disease progression. Funct Integr Genomics. 2021; 21: 451–472. 10.1007/s10142-021-00789-6.

24. Khatun, M.T., Rana, H.K., Hossain, M.A. et al. Bioinformatics and systems biology approaches to identify molecular targets and pathways shared between Schizophrenia and bipolar disorder. Informatics in Medicine Unlocked. 2024; 49: 101556. https://doi.org/10.1016/j.imu.2024.101556.

25. Li, X., Zhuo, S., Cho, Y.S. et al. YAP antagonizes TEAD-mediated AR signaling and prostate cancer growth. EMBO J. 2023; 42: e112184. 10.15252/embj.2022112184.

26. Samarzija, I. Site-Specific and Common Prostate Cancer Metastasis Genes as Suggested by Meta-Analysis of Gene Expression Data. Life (Basel). 2021; 11. 10.3390/life11070636.

27. Xiong, J., Zhang, J. and Li, H. Identification of G2 and S Phase-Expressed-1 as a Potential Biomarker in Patients with Prostate Cancer. Cancer Manag Res. 2020; 12: 9259–9269. 10.2147/CMAR.S272795.

28. Lai, W., Zhu, W., Li, X. et al. GTSE1 promotes prostate cancer cell proliferation via the SP1/FOXM1 signaling pathway. Lab Invest. 2021; 101: 554–563. 10.1038/s41374-020-00510-4.

29. Wang, Y., Wang, J., Yan, K. et al. Identification of core genes associated with prostate cancer progression and outcome via bioinformatics analysis in multiple databases. PeerJ. 2020; 8: e8786. 10.7717/peerj.8786.

30. Patil, M., Pabla, N. and Dong, Z. Checkpoint kinase 1 in DNA damage response and cell cycle regulation. Cell Mol Life Sci. 2013; 70: 4009–4021. 10.1007/s00018-013-1307-3.

31. Drapela, S., Khirsariya, P., van Weerden, W.M. et al. The CHK1 inhibitor MU380 significantly increases the sensitivity of human docetaxel-resistant prostate cancer cells to gemcitabine through the induction of mitotic catastrophe. Mol Oncol. 2020; 14: 2487–2503. 10.1002/1878-0261.12756.

32. Chen, H., Zhou, L., Wu, X. et al. The PI3K/AKT pathway in the pathogenesis of prostate cancer. Front Biosci (Landmark Ed). 2016; 21: 1084–1091. 10.2741/4443.

33. Georgescu, M.M. PTEN Tumor Suppressor Network in PI3K-Akt Pathway Control. Genes Cancer. 2010; 1: 1170–1177. 10.1177/1947601911407325.

34. Lu, D., Song, Y., Yu, Y. et al. KAT2A-mediated AR translocation into nucleus promotes abiraterone-resistance in castration-resistant prostate cancer. Cell Death Dis. 2021; 12: 787. 10.1038/s41419-021-04077-w.

35. Lin, X., Xiang, X., Hao, L. et al. The role of Aurora-A in human cancers and future therapeutics. Am J Cancer Res. 2020; 10: 2705–2729.

36. Kivinummi, K., Urbanucci, A., Leinonen, K. et al. The expression of AURKA is androgen regulated in castration-resistant prostate cancer. Sci Rep. 2017; 7: 17978. 10.1038/s41598-017-18210-3.

37. Chen, X., Ma, J., Wang, X. et al. CCNB1 and AURKA are critical genes for prostate cancer progression and castration-resistant prostate cancer resistant to vinblastine. Front Endocrinol (Lausanne). 2022; 13: 1106175. 10.3389/fendo.2022.1106175.

38. Tooley, J. and Stukenberg, P.T. The Ndc80 complex: integrating the kinetochore’s many movements. Chromosome Res. 2011; 19: 377–391. 10.1007/s10577-010-9180-5.

39. Deng, Y., Li, J., Zhang, Y. et al. NUF2 Promotes Breast Cancer Development as a New Tumor Stem Cell Indicator. Int J Mol Sci. 2023; 24. 10.3390/ijms24044226.

40. Jiang, F., Huang, X., Yang, X. et al. NUF2 Expression Promotes Lung Adenocarcinoma Progression and Is Associated With Poor Prognosis. Front Oncol. 2022; 12: 795971. 10.3389/fonc.2022.795971.

41. Zheng, B., Wang, S., Yuan, X. et al. NUF2 is correlated with a poor prognosis and immune infiltration in clear cell renal cell carcinoma. BMC Urol. 2023; 23: 82. 10.1186/s12894-023-01258-x.

42. Liu, G., Zhu, M., Zhang, M. et al. Emerging Role of IGF-1 in Prostate Cancer: A Promising Biomarker and Therapeutic Target. Cancers (Basel). 2023; 15. 10.3390/cancers15041287.

43. Zhang, Y., Chen, X.N., Zhang, H. et al. CDK13 promotes lipid deposition and prostate cancer progression by stimulating NSUN5-mediated m5C modification of ACC1 mRNA. Cell Death Differ. 2023; 30: 2462–2476. 10.1038/s41418-023-01223-z.

44. Laramas, M., Pasquier, D., Filhol, O. et al. Nuclear localization of protein kinase CK2 catalytic subunit (CK2alpha) is associated with poor prognostic factors in human prostate cancer. Eur J Cancer. 2007; 43: 928–934. 10.1016/j.ejca.2006.11.021.

45. Nassar, Z.D., Mah, C.Y., Dehairs, J. et al. Human DECR1 is an androgen-repressed survival factor that regulates PUFA oxidation to protect prostate tumor cells from ferroptosis. Elife. 2020; 9. 10.7554/eLife.54166.

46. Deshmukh, D. and Qiu, Y. Role of PARP-1 in prostate cancer. Am J Clin Exp Urol. 2015; 3: 1–12.

47. Deng, X., Shao, G., Zhang, H.T. et al. Protein arginine methyltransferase 5 functions as an epigenetic activator of the androgen receptor to promote prostate cancer cell growth. Oncogene. 2017; 36: 1223–1231. 10.1038/onc.2016.287.

48. Owens, J.L., Beketova, E., Liu, S. et al. Targeting Protein Arginine Methyltransferase 5 Suppresses Radiation-induced Neuroendocrine Differentiation and Sensitizes Prostate Cancer Cells to Radiation. Mol Cancer Ther. 2022; 21: 448–459. 10.1158/1535-7163.MCT-21-0103.

49. Ngo, V.N., Davis, R.E., Lamy, L. et al. A loss-of-function RNA interference screen for molecular targets in cancer. Nature. 2006; 441: 106–110. 10.1038/nature04687.

50. Hart, T., Chandrashekhar, M., Aregger, M. et al. High-Resolution CRISPR Screens Reveal Fitness Genes and Genotype-Specific Cancer Liabilities. Cell. 2015; 163: 1515–1526. 10.1016/j.cell.2015.11.015.

51. Yang, X., Liu, J., Wang, S. et al. Genome wide-scale CRISPR-Cas9 knockout screens identify a fitness score for optimized risk stratification in colorectal cancer. J Transl Med. 2024; 22: 554. 10.1186/s12967-024-05323-3.

52. Viswanatha, R., Li, Z., Hu, Y. et al. Pooled genome-wide CRISPR screening for basal and context-specific fitness gene essentiality in Drosophila cells. Elife. 2018; 7. 10.7554/eLife.36333.

53. Wei, L., Lee, D., Law, C.T. et al. Genome-wide CRISPR/Cas9 library screening identified PHGDH as a critical driver for Sorafenib resistance in HCC. Nat Commun. 2019; 10: 4681. 10.1038/s41467-019-12606-7.

